# Clinical, imaging genetics and deformation based morphometry study of longitudinal changes after surgery for intractable aggressive behaviour

**DOI:** 10.1101/548826

**Authors:** Flavia V. Gouveia, Jürgen Germann, Rosa M. C. B. de Morais, Erich T. Fonoff, Clement Hamani, Eduardo Joaquim Alho, Helena Brentani, Ana Paula Martins, Gabriel Devenyi, Raihaan Patel, Christopher J. Steele, Robert Gramer, M. Mallar Chakravarty, Raquel C. R. Martinez

**Affiliations:** Laboratory of Neuromodulation, Teaching and Research Institute, Hospital Sirio-Libanes, Sao Paulo, Sao Paulo, Brazil; Neuromodulation Centre, Sunnybrook Reseach Institute, University of Toronto, Toronto, Ontario, Canada; CIC, Douglas Mental Health University Institute, McGill University, Montreal, Quebec, Canada; PROTEA, Department of Psychiatry - University of Sao Paulo School of Medicine, Sao Paulo, Sao Paulo, Brazil; Department of Neurology, Division of Functional Neurosurgery of the Institute of Psychiatry, University of Sao Paulo, Medical School, Sao Paulo, Sao Paulo, Brazil; Division of Neurosurgery, Sunnybrook Health Sciences Centre, University of Toronto, Toronto, Ontario, Canada; Department of Psychiatry - University of Sao Paulo, Medical School, Sao Paulo, Sao Paulo, Brazil; National Institute of Developmental Psychiatry for Children and Adolescents, CNPq, Sao Paulo, Sao Paulo, Brazil; Department of Neurosurgery, Duke University Medical Center, Durham, North Carolina, United States

**Keywords:** Aggressive Behavior, Amygdala, Testosterone, Magnetic Ressonance Imaging, Deformation Based Morphometry, Neurosurgery

## Abstract

Intractable aggressive behaviour is a devastating behavioural disorder that reach 30% of psychiatric aggressive patients. Neuromodulatory surgeries may be treatment alternatives to reduce suffering. We investigated the outcomes of bilateral amygdala radiofrequency ablation in four patients with intractable aggressive behaviour (life-threatening-self-injury and social aggression) by studying whole brain magnetic resonance imaging and clinical data. Post-surgery assessments revealed decreases in aggression and agitation and improvements in quality of life. Aggressive behaviour was positively correlated with serum testosterone levels and the testosterone/cortisol ratio in males. No clinically significant side effects were observed. Imaging analyses revealed preoperative amygdala volumes within normal range and confirmed appropriate lesion locations. Reduction in aggressiveness were accompanied by volumetric reduction in brain areas associated with aggressive behaviour (express genes related to aggressive behaviour), and increases in regions related to somatosensation. These findings further elucidates the neurocircuitry of aggression and suggests novel neuromodulation targets.

## INTRODUCTION

Excessive aggressive behavior is highly prevalent in patients with Autism Spectrum Disorder (ASD) and Intellectual Disabilities (ID)^1^. It presents a major obstacle to patient care and treatment^2^. Additionally, it often presents as severe self-injurious behavior that may culminate with the need for chronic restraining of patients, substantially reducing quality of life^3,4^. The primary treatment for excessive aggressive behavior involves the use of behavioral therapy and medications that act mainly on dopaminergic and serotoninergic systems^1,5^. When patients fail to respond to conventional behavioral and medical treatments, high dose mono- or polypharmacy strategies are indicated^6^. Although these therapies are successful in most patients, there is a significant subset of individuals (approximately 30%) who do not adequately respond to treatment and are considered medication-refractory^7^.

Structures related to the control of aggressive behavior include the prefrontal cortex, amygdala, hypothalamus and midbrain periaqueductal gray^8^. It is believed that aggressive individuals have alterations in these structures. Prior research suggests that higher limbic system activity alters perception of provocative stimuli, and reduces “top-down” control of the prefrontal cortex, minimizing the ability to suppress aggressive behavior^8^. Lesion studies performed in the late 18^th^ and early 19^th^ centuries have shown that the removal of the temporal lobes or selective amygdala ablation resulted in marked reduction of aggressive behavior in humans^9–11^ and nonhuman primates^12,13^. Imaging studies have been largely controversial and/or inconclusive in asserting an association between amygdala volume^14^ or activity^15^ and behavioral disorders (including aggressive behavior). For example, veterans with aggressive behavior disorders have been shown to demonstrate a more intense amygdala response to external stimuli as well as an attenuated amygdala-prefrontal cortex connectivity^15,16^. In contrast, studies in criminal psychopaths have reported that the level of amygdala activation is lower during processing of negative affective stimuli, fear conditioning paradigms, and emotional moral decision making^17,18^.

More recently, amygdala deep brain stimulation (DBS) was successfully used to reduce aggressive behavior (life-threatening self-injurious behavior) in a drug-refractory autistic teenager^3^. Quite often, however, patients with severe self-aggressive behavior towards the face and head are precluded from DBS consideration for fear of complications such as infection, skin erosion, and lead fracture^19^. For this small population of non-responsive individuals, stereotactic ablative surgery targeting the amygdala has been performed with beneficial result^11,20^. As this patient population is small and the number of recent publications are limited, in depth clinical follow up and the study of neuroanatomical and/or functional aspects of this surgery remain sparse. In particular, to date, no study has attempted to identify and characterize the brain morphology and changes associated with surgical intervention in these subjects.

We addressed the crucial gap in understanding the aggressive brain circuitry and how structural and functional changes in this network relate to behavior changes by studying longitudinal local volume and structure changes throughout the whole brain and examining the clinical and hormonal effects of bilateral amygdala radiofrequency ablation in a small cohort of patients with aggressive behavior.

## RESULTS

Behavior assessments demonstrated a marked reduction in aggressive behavior in all patients (p01: 84.6%; p02: 80%; p03: 88.9%; p04: 56.2%; p_(cor)_<0.01; Figure 1E). This was associated with a reduction in motor agitation (p_(cor)_<0.05; Figure 1F) and an increase in quality of life (p_(cor)_<0.01; Figure 1G). Significant correlations were found between improvements in aggressive behavior and agitation (p<0.0001) and between improvements in aggressive behavior and quality of life (p=0.01612; Supplementary Figures 1A-B).

**Figure 1.**
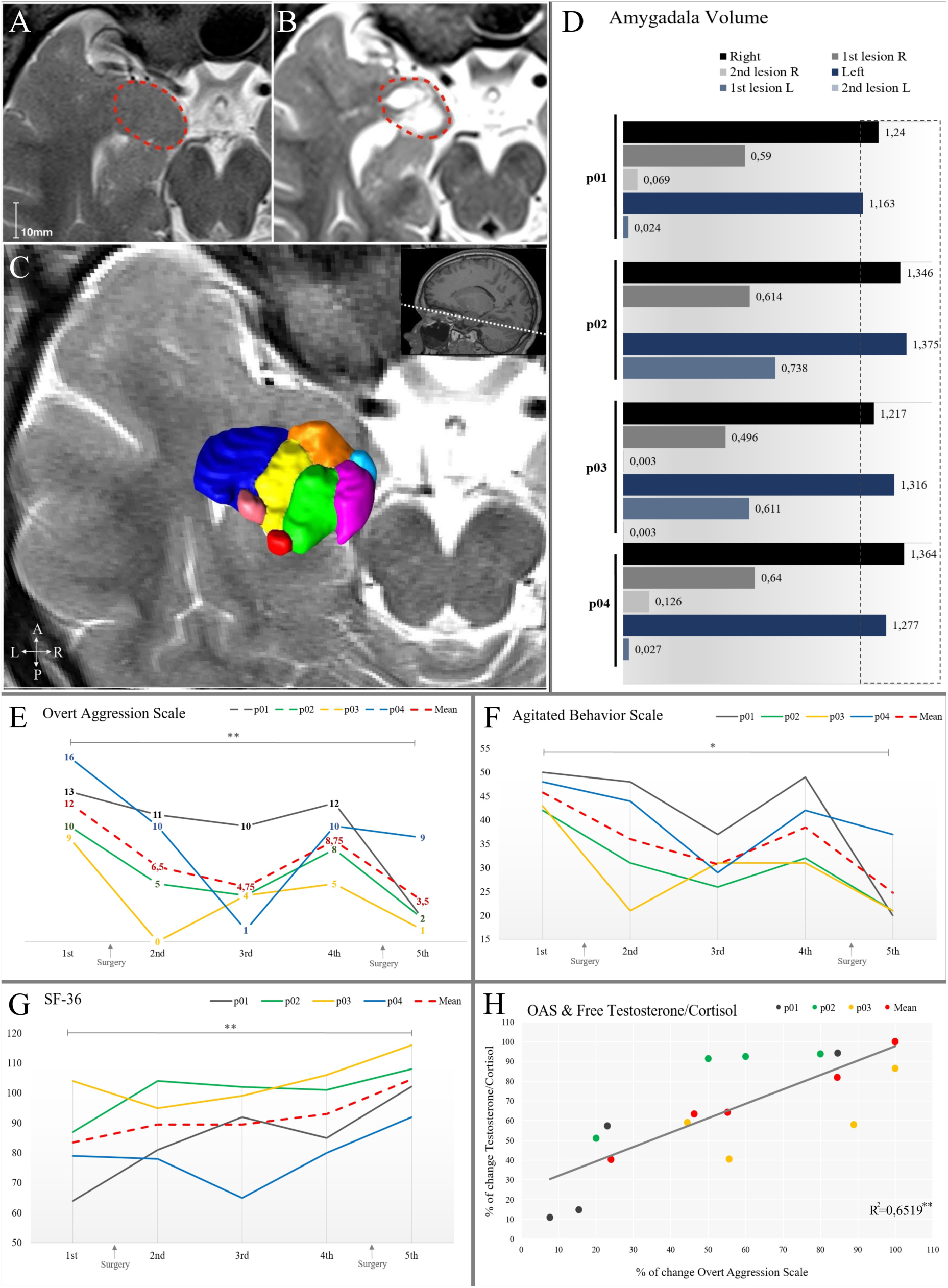
Baseline and follow up evaluations. A. MRI image showing amygdala location before surgery. B. MRI image showing lesion placement. C. 3D reconstruction of the amygdala on an MRI image. D. Amygdala volume; E. Aggressive levels measured by the Overt Aggression Scale (OAS); F. General motor agitation measured by the Agitated Behavior Scale (ABS); G. Quality of life evaluated by the Short Form Survey (SF-36); H: Correlation between the testosterone/cortisol ratio and aggressive behavior for male patients (p01, p02, p03, mean). Abbreviations: 1st: Baseline evaluation; 2nd: 3 months after surgery; 3rd: 12 months after surgery; 4th: 24 months after surgery; 5th: 36 months after surgery for p03 and after 2nd surgery for p01, p02 and p04. *p<0.05; **p<0.01.

Each total amygdala volume was measured before the surgical procedure and was determined to be within normal range^21^ (Figure 1A-D). After surgery, we confirmed the location of the lesion and measured its size. As shown in Figure 1D, lesions were smaller in volume than the amygdala. P01, p02 and p04 required a second procedure, after which most of the amygdala was ablated.

The ratio between testosterone and cortisol is a biomarker for aggressive behavior^22^. We identified a positive correlation between aggressive behavior and the testosterone/cortisol ratio in the males (p_(cor)_<0.01, Figure 1H; p01 p=0.0205; p02 p=0.0243; p03 p=0.0334; p04 p=0.315; Supplementary Figures 1D-G) and between aggressive behavior and total testosterone, also only in the male patients (p=0.00101; p01p=0.0142; p02 p=0.0584; p03 p=0.0247; p04 p=0.8059, Supplementary Figures 1C and 2H-K). Although some fluctuation in the hormone levels occurred, there was no significant clinical impact on patients well-being (Supplementary Table 1).

**Table 1.**
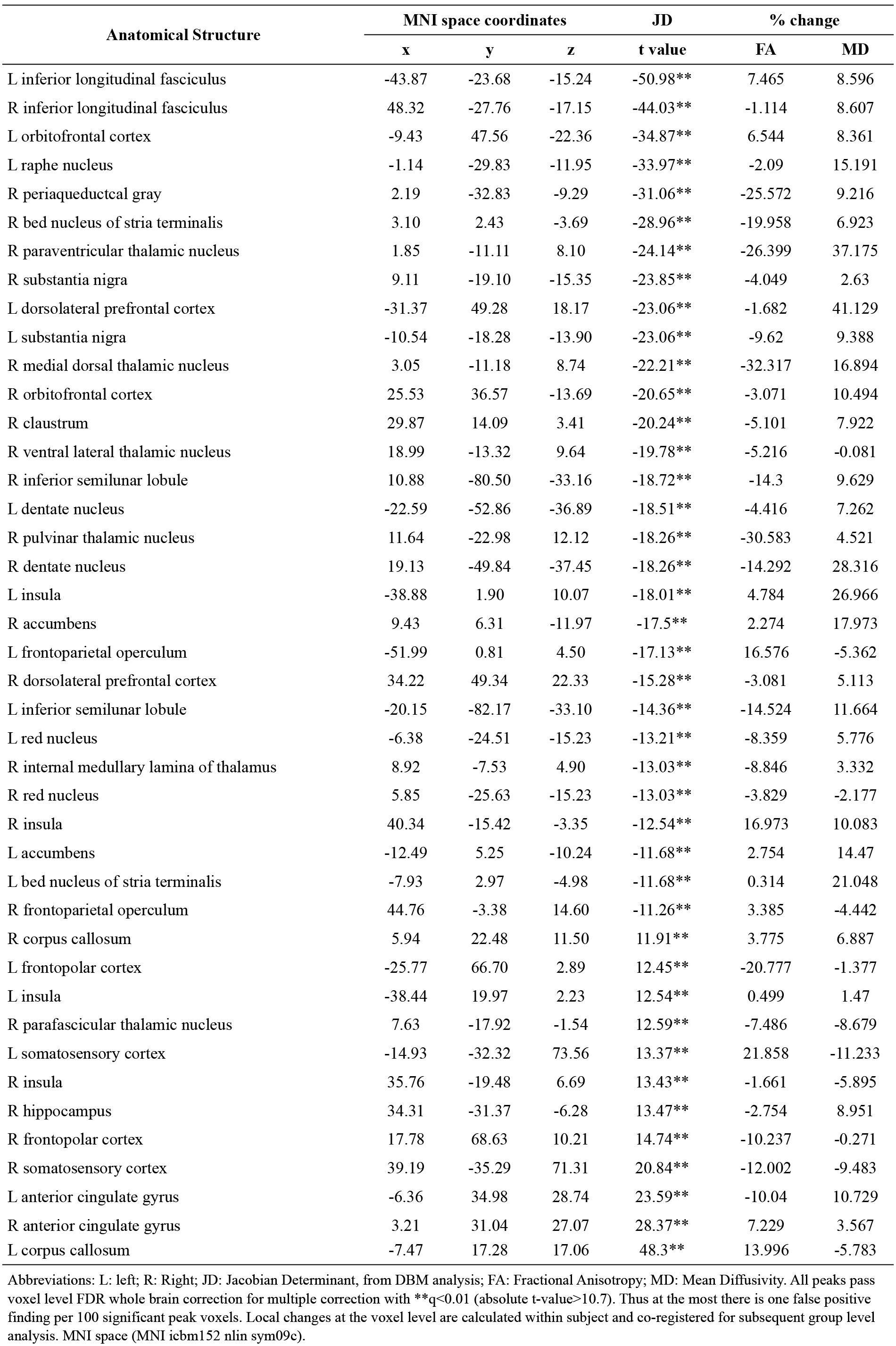
Longntudinal analysis with DBM and FA/MD maps

| Anatomical Structure | MNI space coordinates |  |  | JD | % change |  |
| --- | --- | --- | --- | --- | --- | --- |
|  | x | y | z | t value | FA | MD |
| L inferior longitudinal fasciculus | -43.87 | -23.68 | -15.24 | -50.98** | 7.465 | 8.596 |
| R inferior longitudinal fasciculus | 48.32 | -27.76 | -17.15 | -44.03** | -1.114 | 8.607 |
| L orbitofrontal cortex | -9.43 | 47.56 | -22.36 | -34.87** | 6.544 | 8.361 |
| L raphe nucleus | -1.14 | -29.83 | -11.95 | -33.97** | -2.09 | 15.191 |
| R periaqueductal gray | 2.19 | -32.83 | -9.29 | -31.06** | -25.572 | 9.216 |
| R bed nucleus of stria terminalis | 3.10 | 2.43 | -3.69 | -28.96** | -19.958 | 6.923 |
| R paraventricular thalamic nucleus | 1.85 | -11.11 | 8.10 | -24.14** | -26.399 | 37.175 |
| R substantia nigra | 9.11 | -19.10 | -15.35 | -23.85** | -4.049 | 2.63 |
| L dorsolateral prefrontal cortex | -31.37 | 49.28 | 18.17 | -23.06** | -1.682 | 41.129 |
| L substantia nigra | -10.54 | -18.28 | -13.90 | -23.06** | -9.62 | 9.388 |
| R medial dorsal thalamic nucleus | 3.05 | -11.18 | 8.74 | -22.21** | -32.317 | 16.894 |
| R orbitofrontal cortex | 25.53 | 36.57 | -13.69 | -20.65** | -3.071 | 10.494 |
| R claustrum | 29.87 | 14.09 | 3.41 | -20.24** | -5.101 | 7.922 |
| R ventral lateral thalamic nucleus | 18.99 | -13.32 | 9.64 | -19.78** | -5.216 | -0.081 |
| R inferior semilunar lobule | 10.88 | -80.50 | -33.16 | -18.72** | -14.3 | 9.629 |
| L dentate nucleus | -22.59 | -52.86 | -36.89 | -18.51** | -4.416 | 7.262 |
| R pulvinar thalamic nucleus | 11.64 | -22.98 | 12.12 | -18.26** | -30.583 | 4.521 |
| R dentate nucleus | 19.13 | -49.84 | -37.45 | -18.26** | -14.292 | 28.316 |
| L insula | -38.88 | 1.90 | 10.07 | -18.01** | 4.784 | 26.966 |
| R accumbens | 9.43 | 6.31 | -11.97 | -17.5** | 2.274 | 17.973 |
| L frontoparietal operculum | -51.99 | 0.81 | 4.50 | -17.13** | 16.576 | -5.362 |
| R dorsolateral prefrontal cortex | 34.22 | 49.34 | 22.33 | -15.28** | -3.081 | 5.113 |
| L inferior semilunar lobule | -20.15 | -82.17 | -33.10 | -14.36** | -14.524 | 11.664 |
| L red nucleus | -6.38 | -24.51 | -15.23 | -13.21** | -8.359 | 5.776 |
| R internal medullary lamina of thalamus | 8.92 | -7.53 | 4.90 | -13.03** | -8.846 | 3.332 |
| R red nucleus | 5.85 | -25.63 | -15.23 | -13.03** | -3.829 | -2.177 |
| R insula | 40.34 | -15.42 | -3.35 | -12.54** | 16.973 | 10.083 |
| L accumbens | -12.49 | 5.25 | -10.24 | -11.68** | 2.754 | 14.47 |
| L bed nucleus of stria terminalis | -7.93 | 2.97 | -4.98 | -11.68** | 0.314 | 21.048 |
| R frontoparietal operculum | 44.76 | -3.38 | 14.60 | -11.26** | 3.385 | -4.442 |
| R corpus callosum | 5.94 | 22.48 | 11.50 | 11.91** | 3.775 | 6.887 |
| L frontopolar cortex | -25.77 | 66.70 | 2.89 | 12.45** | -20.777 | -1.377 |
| L insula | -38.44 | 19.97 | 2.23 | 12.54** | 0.499 | 1.47 |
| R parafascicular thalamic nucleus | 7.63 | -17.92 | -1.54 | 12.59** | -7.486 | -8.679 |
| L somatosensory cortex | -14.93 | -32.32 | 73.56 | 13.37** | 21.858 | -11.233 |
| R insula | 35.76 | -19.48 | 6.69 | 13.43** | -1.661 | -5.895 |
| R hippocampus | 34.31 | -31.37 | -6.28 | 13.47** | -2.754 | 8.951 |
| R frontopolar cortex | 17.78 | 68.63 | 10.21 | 14.74** | -10.237 | -0.271 |
| R somatosensory cortex | 39.19 | -35.29 | 71.31 | 20.84** | -12.002 | -9.483 |
| L anterior cingulate gyrus | -6.36 | 34.98 | 28.74 | 23.59** | -10.04 | 10.729 |
| R anterior cingulate gyrus | 3.21 | 31.04 | 27.07 | 28.37** | 7.229 | 3.567 |
| L corpus callosum | -7.47 | 17.28 | 17.06 | 48.3** | 13.996 | -5.783 |

Abbreviations: L: left; R: Right; JD: Jacobian Determinant, from DBM analysis; FA: Fractional Anisotropy; MD: Mean Diffusivity. All peaks pass voxel level FDR whole brain correction for multiple correction with $**q < 0.01$ (absolute t-value $> 10.7$ ). Thus at the most there is one false positive finding per 100 significant peak voxels. Local changes at the voxel level are calculated within subject and co-registered for subsequent group level analysis. MNI space (MNI icbm152 nlin sym09c).

The longitudinal DBM analysis of these patients (Figure 2) identified significant morphological changes in several brain regions. Volumetric decrease was found in many areas, including those known to play a role in mechanisms of aggressive behavior (e.g. doroslateral prefrontal cortex (DLPFC), nucleus accumbens (NAcc), bed nucleus of stria terminalis (BNST))^8,23^. Volume increases were observed in areas involved in somatosensory processing and pain perception (e.g. somatosensory cortex, thalamus)^24^. Supplementary Figure 2 illustrates the volume change in those and other main regions. Table 1 lists and names the key peaks passing FDR correction (q<0.01) their t-value and FA and MD percent change.

**Figure 2.**
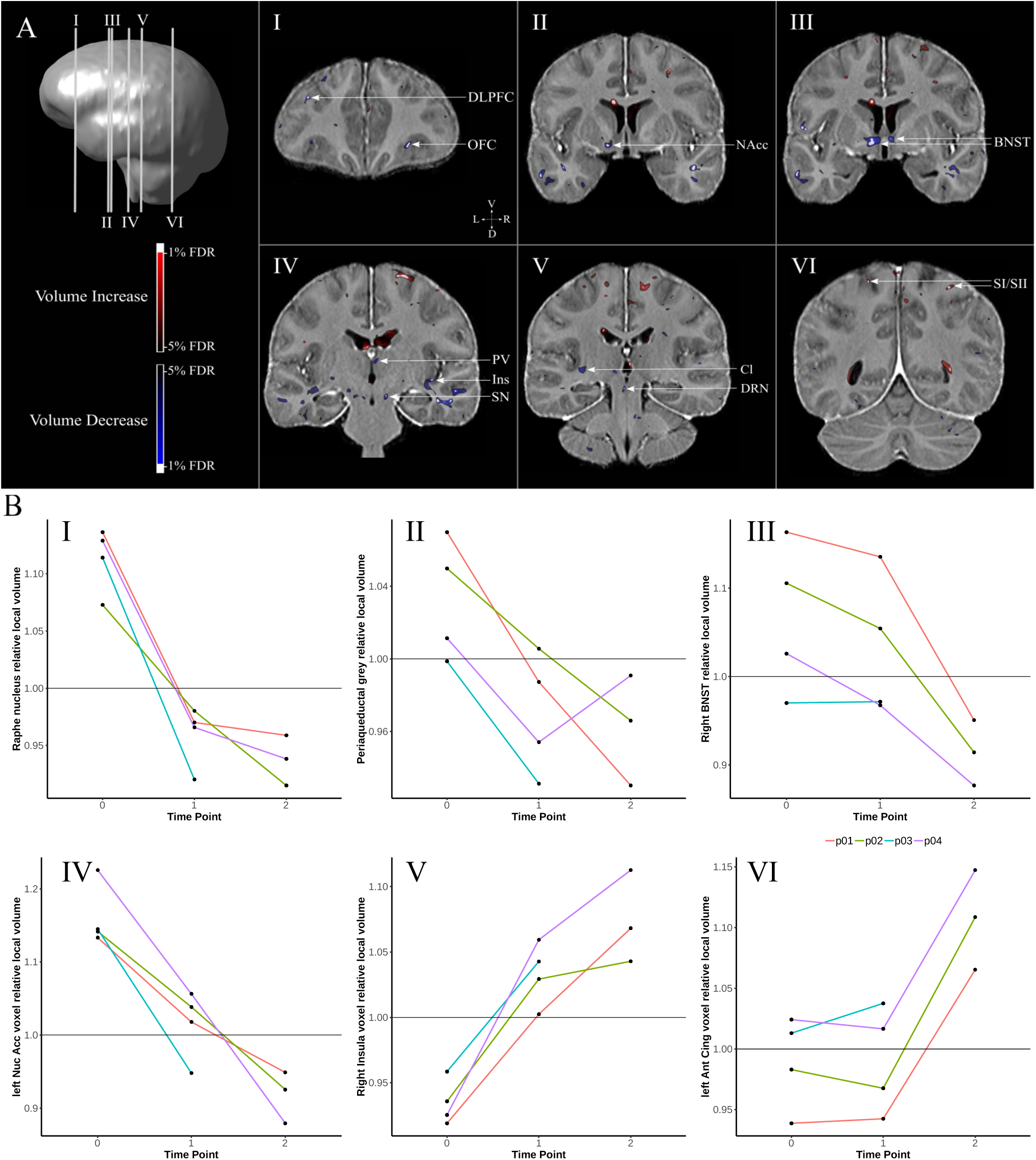
Results from Deformation Based Morphometry analysis (DBM). A. Coronal slices illustrating areas that presented significant local volume changes in the longitudinal DBM analysis. The areas named are known to be part of the neurocircuitry of aggressive behavior or somatosensation. B. Changes overtime in selected peak voxels in each patient. Data is presented as scale of relative voxel size. I. Dorsal Raphe nucleus; II. Periaqueductal Gray; III. Right Bed Nucleus of Stria Terminalis; IV. Left Nucleus Accumbens; V. Right Insula; VI. Left Anterior Cinculate. Abbreviations: BNST: bed nucleus of stria terminalis; Cl: claustrum; cx: cortex; DLPFC: dorsolateral prefrontal cortex; DRN: dorsal raphe nucleus; Ins: insula; int. internal; L: left; lam: lamina; NAcc: nucleus accumbens; nu: nucleus; OFC: orbitofrontal cortex; PV: paraventricular thalamic nucleus; R:right; SI/SII: somatosensory cortex; SN: substantia nigra.

Likewise, both FA and MD changed after surgery and there was a significant correlation between changes in Jacobians and MD, which is related to the displacement of water (FA p_(cor)_>0.05; MD p_(cor)_<0.05; Figure 3A for main peaks. Supplementary Figure 3 for all peaks).

**Figure 3.**
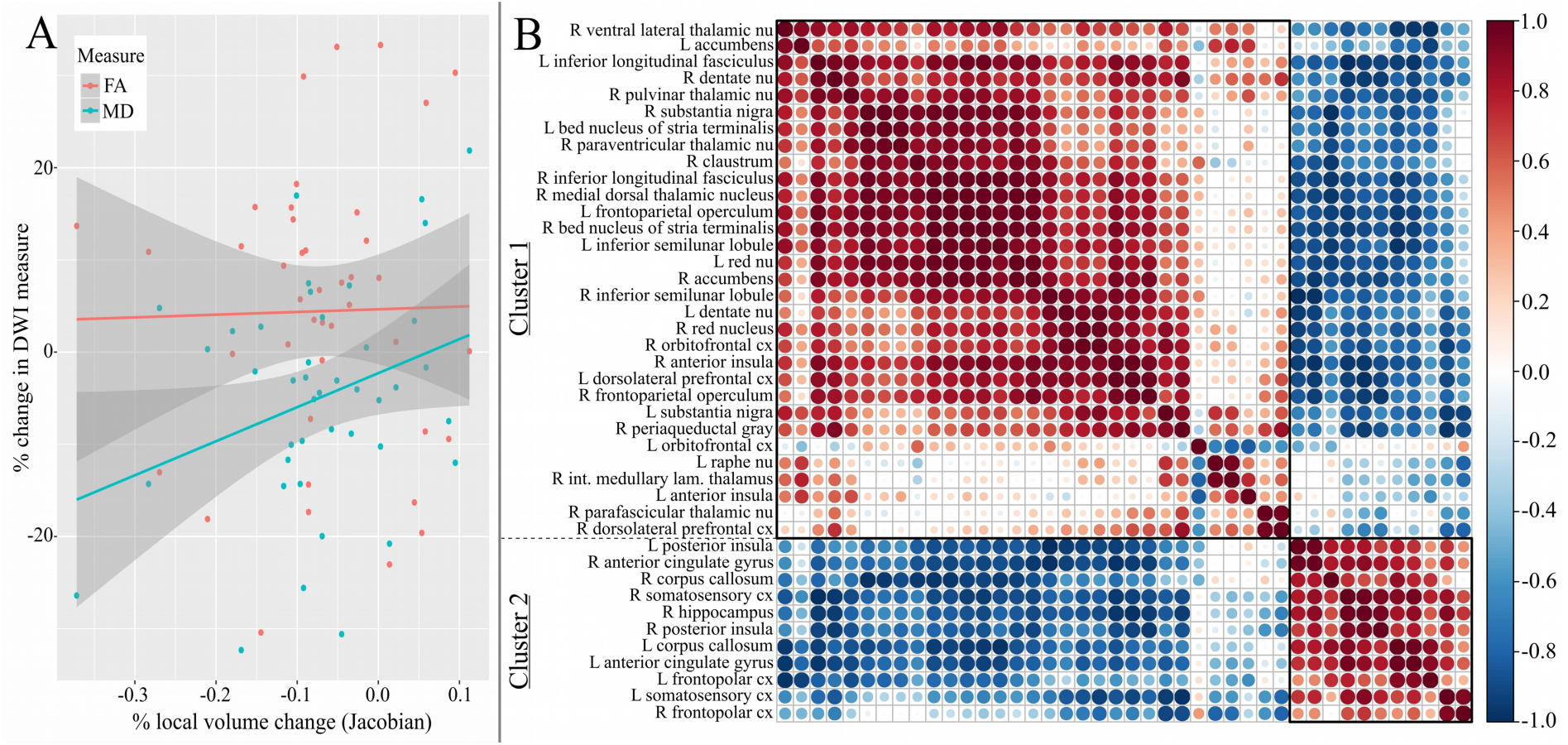
Results from Fractional Anisotropy (FA) and Mean Diffusivity (MD) maps and Cluster analysis of the Deformation Based Morphometry study. A. Correlation between DBM Jacobians, FA and MD maps for main peaks. B. Hierarchical cluster of DBM peaks (5% FDR correction).

Hierarchical clustering was applied to investigate the possibility of groupings within brain areas that were significant in the DBM analysis. The results confirmed the presence of 2 clusters: Cluster 1: areas that decrease in volume known to be related to aggressive behavior and connected to the amygdala^8,23^; Cluster 2: areas that increase in volume involved in somatosensation and pain perception^24^ (Figure 3B).

The gene ontology analysis of the genes highly expressed in regions that exhibited morphological shrinkage (Cluster 1) compared to the rest of the brain showed functions related to amygdala and aggressive behavior: 1. Steroid metabolic process; 2. Regulation of grooming behavior; 3. Response to chemicals Nitric oxide and Amphetamine; 4. Aminergic neurotransmitter transport. Figure 4 shows these results and the complete gene enrichment analysis can be found in Supplementary Figure 4.

**Figure 4.**
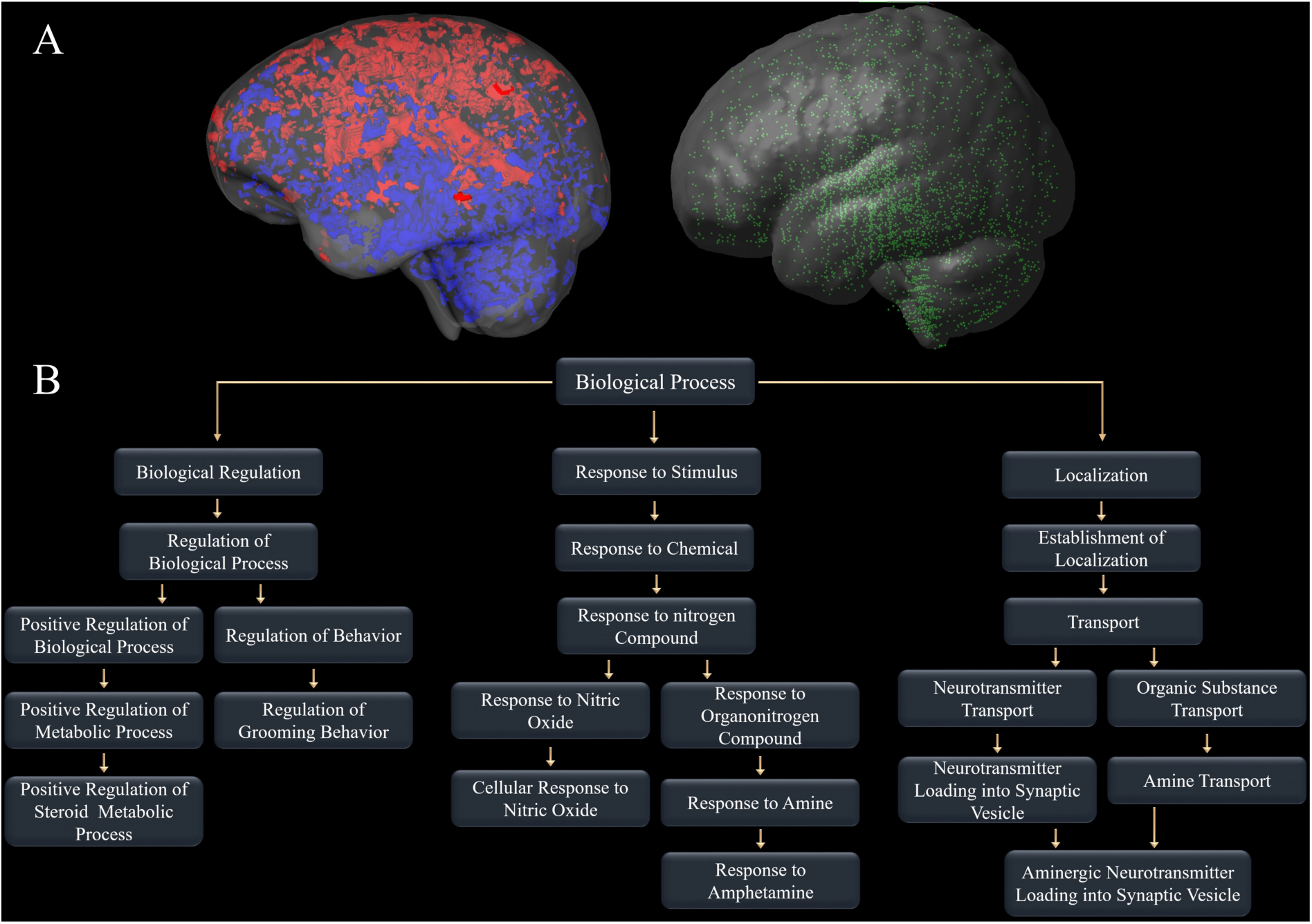
Results of gene enrichment analysis. A. Illustration of the inside/area genetic enrichment analysis. The left side shows the distribution of DBM results (10% FDR corrected) red represents increases and blue decreases in volume. The right shows the location where tissue samples were collected for the gene expression analysis. Every green dot indicates a well location. B. Summary of GOrilla and GEAS gene enrichment analysis results for decreased Jacobians and overexpressed genes.

## DISCUSSION

Stereotactic surgery led to a remarkable reduction in aggressive behavior and improvements in quality of life in patients with medically-refractory aggressive behavior. To assess potential therapeutic mechanisms for the procedure, MRI images acquired before and after the surgery were analyzed using DBM, and FA and MD maps computed. While the DBM technique uses image registration to calculate local deformation fields at a voxel level for the detection of morphological variations^25^, FA and MD values are indicative of local microstructural characteristics primarily related to white matter. FA is an index of diffusion asymmetry and is related to characteristics such as fiber density, axonal diameter and myelination, while MD shows the degree of displacement of water molecules^26^.

We found that morphological changes associated with reduction in aggressive behavior could be grouped into two clusters: Cluster 1 includes areas associated with aggressive behavior such as the DLPFC, NAcc, BNST, Insula, periaqueductal gray (PAG), and dorsal raphe nucleus (DRN). Cluster 2 regions includes areas related to somatosensation and pain perception, such as somatosensory cortex, thalamus, and anterior Cingulate Cortex. The correlation of volumetric changes with DWI measures suggests that the volume change following surgery is related to local microstructural changes. Though the exact mechanism driving these distinct changes remains unexplained (e.g. increase/decrease of number of glial cells, hypo- or hypertrophy of neurons), it is interesting to note the clusters change in opposite directions. Areas of cluster 1 receive alert signals from the presumably hyperactive amygdala^8^ and decrease in volume once the amygdala stimulation ceases. Whereas, activity in the regions of cluster 2 are suppressed when the amygdala is highly activated and increase in volume post-surgery. The distinct pattern of local changes throughout the somatosensory/pain network is particularly intriguing in this group and may explain the reduction in self-aggressive behavior observed after surgery (OAS self-injury subscale change: p01: 4 to 1; p02:1 to 0; p03: 4 to 0; p04: 4 to 2). It is known that the amygdala has a central role in processing nociceptive^27^ stimuli and that patients with self-injurious behavior have altered pain perceptior^28^. Thus, amygdala ablation could affect this neurocircuitry, restoring normal activity in areas related to the process of nociceptive stimuli resulting in reduction of self-injury behavior.

The correlation found between aggressive behavior and agitation supports the theory that excessive aggressive behavior occurs when the amygdala is hyper-activated and more responsive to environmental stimuli^8^. In fact, after surgery, patients were able to return to a special needs school that they were previously forced to withdraw from when their condition had progressively worsened. There they are able to learn and improve skills, mainly related to daily life activity and communication. In this sense, the procedure appears to leads to significant behavioral improvement which enables patients to function in a social construct that fosters further behavioral improvement. Although permanent side effects and worsened behavioral problems have been previously reported^9,11,19^, an important aspect of our study was that no noticeable side effects were observed after surgery.

The gene ontology analysis found that Cluster 1 regions are related to steroid metabolic process, response to Nitric oxide and Amphetamine, aminergic neurotransmitter transport, and interestingly, regulation of grooming behavior. These all are closely related to the amygdala and aggressive behavior: Grooming behavior is similar to aggressive behaviors in the sense that it is an innate social behavior influenced by the amygdala, which plays an important role in social bonding and the maintenance of species^29^. The aminergic neurotransmitter system (e.g. serotonin and dopamine) as well as nitric oxide and amphetamine have all been implicated in the modulation of aggressive behavio^5,30,31^The mechanism for this effect, however, is still unknown. Steroid metabolism is a process linked to aggressive behavior and is influenced by the amygdala via extensive bi-directional projections to and from the hypothalamus. This latter structure is responsible for controlling hormonal production and secretion including precursors of the synthesis of testosterone and cortisol^32^. These two hormones are part of internal feedback loops of the amygdala resulting in the expression of more aggressive actions (consequence of testosterone augmentation) or increased fear responses (consequence of cortisol predominance). It is believed that the ratio between those two hormones may be a biomarker for aggressive behavior^22^. In fact, this is reinforced by the correlation found between testosterone and the testosterone/cortisol ratio and aggression.

To our knowledge, this is the first report of longitudinal neuroimaging analyses of patients treated with bilateral amygdala radiofrequency ablation. DBM analyses enabled the detection of subtle local volumetric changes throughout the brain and helped us identify numerous local structural alterations that accompany the behavioral changes. We observed consistent characteristic changes within our patients and we believe that this analysis better elucidates the whole human neurocircuitry of aggressive behavior and provides new insights for future interventions. All the changes result in striking behavior alterations leading to easing of suffering and improvement in quality of life. The changes essential and necessary for the positive outcome are yet to be elucidated. Expounding these connections through future research could be instrumental in deriving a better understanding this complex neurocircuitry as well as its mechanism in modulating behavior. Our results, however, suggest an interesting starting point to investigate possible new targets for intervention. For example, one could propose the use of non-invasive neuromodulatory techniques (e.g. Transcranial Magnetic Stimulation (TMS), Transcranial direct current stimulation (tDCS)) delivered to somatosensory cortical areas or DBS to the NAcc or BNST, for the treatment of patients with excessive aggressive behavior. Furthermore, these results highlight the need for studies to ascertain the cause of morphological changes, identify the population of neurons being affected, and explore the impact of these changes on neurotransmitter systems, especially in the context of the aminergic system.

In summary, our results provide novel evidence that bilateral amygdala ablation induces re-adaptation of the neurocircuitry that controls aggressive behavior, with a concomitant behavioral phenotype. These morphological changes could be integral to the reduction of aggressive behavior and increase in quality of life observed in our study population. Further research is necessary to better understand this complex neurocircuitry and explore new targets for therapeutic interventions and less invasive techniques.

## METHODS AND MATERIALS

### SUBJECTS

Four patients with intractable aggressive behavior (3 males, 1 female; aged 19-32 years; demographics in Table 2) were included in the study following psychiatric referral from our outpatient psychiatric clinic. All subjects were diagnosed with severe ID and the 3 males also with ASD. All patients showed uncontrollable life-threatening-self-injurious behavior and extreme aggressive behavior towards their surroundings and others. Three patients required restraint measures on a daily basis. The subjects were selected for the ablation procedure as their severe self-aggressive behavior made them non-eligible for DBS^19.^ An experienced multidisciplinary team assisted the patients and caregivers at all times. Patients’ psychiatric histories are presented in Supplementary Text 1. Criteria for patient selection and for defining treatment refractoriness followed the mandate for conducting psychiatric surgery stipulated by the Regional and Federal Medical Councils. All study procedures, logs, and results were documented and archived in accordance with the Research Ethical Board (#CAPPesq742.331; #CAAE31828014.6.3001.5461). The patient’s parents provided written informed consent after being instructed about potential risks and benefits of surgical treatment. The study was registered as a clinical trial at https://clinicaltrials.gov (#NCT03452878).

**Table 2.** Demographics and clinical characteristics

| Patient | Karyotype | Main diagnosis | Intellectual capacity | Preoperative medication |
| --- | --- | --- | --- | --- |
| p01 | 46,XY normal | Autism spectrum disorder | Severely impaired | Aripiprazole; Biperiden; Carbamazepine; Chlorpromazine; Clozapine; Divalproex sodium; Haloperidol; Levomepromazine; Olanzapine; Promethazine; Quetiapine fumarate; Risperidone; Thioridazine; Ziprasidone |
| p02 | 46,XY normal | Autism spectrum disorder | Severely impaired | Aripiprazole; Benzodiazepines; Biperiden; Carbamazepine; Chlorpromazine; Clonidine; Clozapine; Divalproex sodium; Haloperidol; Levomepromazine; Olanzapine; Promethazine; Risperidone; Thioridazine; Topiramate |
| p03 | 46,XY<br>normal | Autism<br>spectrum<br>disorder | Severely<br>impaired | Aripiprazole; Biperiden; Carbamazepine;<br>Chlorpromazine; Clomipramine; Divalproex sodium;<br>Haloperidol; Levomepromazine; Olanzapine;<br>Oxcarbazepine; Promethazine; Quetiapine fumarate;<br>Risperidone; Sertraline; Thioridazine; Topiramate;<br>Ziprasidone |
| p04 | 46,XX<br>normal | Intellectual<br>disabilities | Severely<br>impaired | Aripiprazole; Carbamazepine; Chlorpromazine;<br>Divalproex sodium; Fluoxetine; Haloperidol;<br>Levomepromazine; Lithium; Methylphenidate;<br>Mirtazapine; Olanzapine; Quetiapine fumarate;<br>Risperidone; Topiramate |

### PRE AND POSTOPERATIVE EVALUATIONS

Preoperative assessment involved full clinical, psychiatric, and neurosurgical evaluations as well as whole brain magnetic resonance imaging (MRI). Questionnaires quantified aggressive behavior, general motor agitation, and quality of life with the Overt Aggression Scale (OAS), Agitated Behavior Scale (ABS), and Short Form Survey (SF-36), respectively. Laboratory tests included: thyroid, kidney, and liver function, adrenal and gonadal hormone levels; lipid, coagulation, and inflammation profiles; and complete blood count (Supplementary Table 1). Since patients presented with severe intellectual disability, neuropsychological evaluations were deferred.

Postoperative follow up included all preoperative measures and was performed at 3, 12, 24 and 36 months after surgery. Patients p01, p02 and p04 needed a second surgery 24 months after the first to increase lesion size, characterized on imaging as well as by minimal symptomatic benefits. Their follow-up was indexed from the date of the first operation. Supplementary Figure 5 illustrates the study timeline.

**Figure 5.**
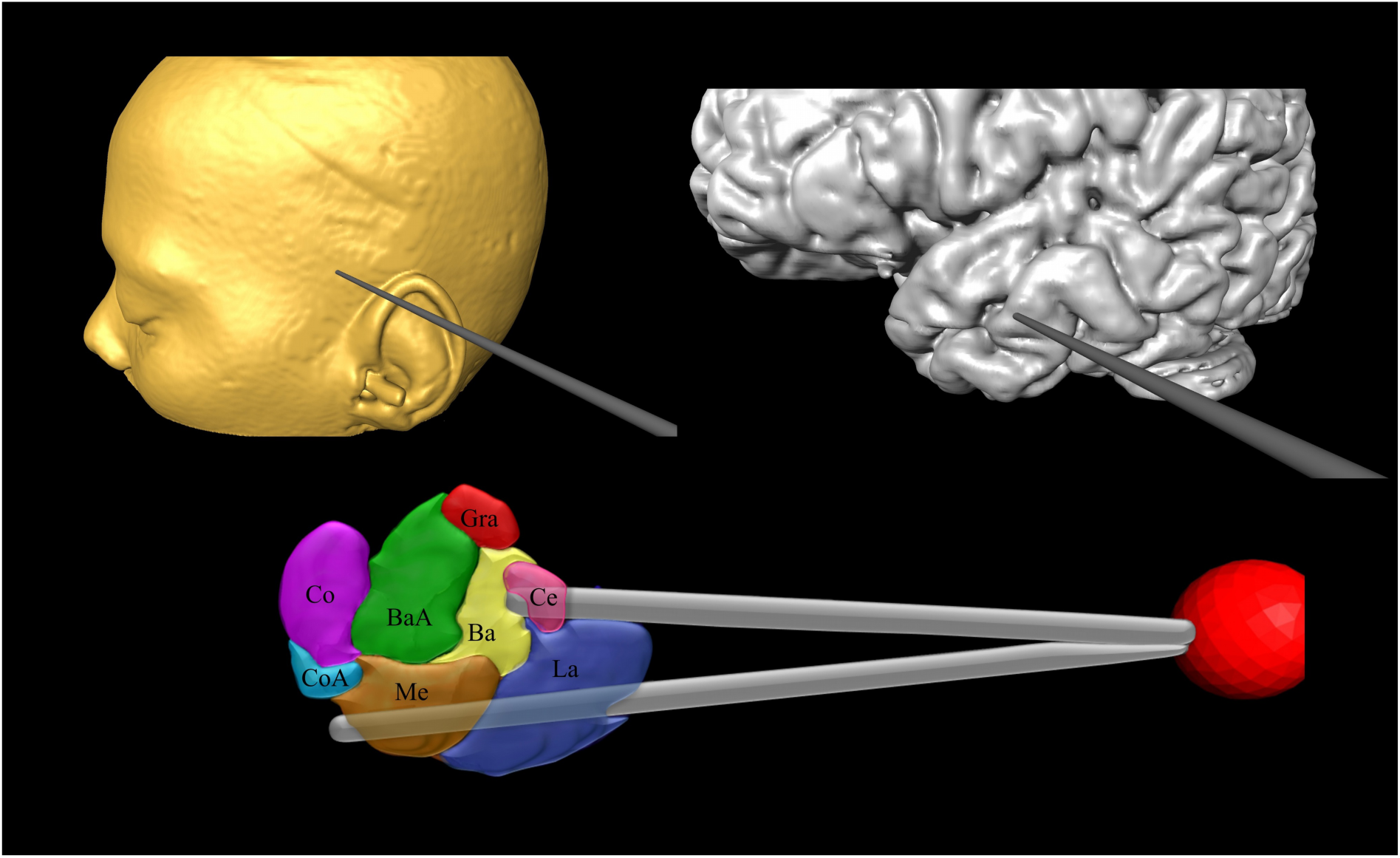
3D reconstructions of probe placement in the amygdala. Abbreviations: Ba: Basalis; BaA: Basalis acessorius; Ce: Centralis; Co: Corticalis; CaA: Corticalis acessorius; Gra: Granularis; La: Lateralis; Me: Medialis.

### SURGERY

The surgeries were completed under local anesthesia and sedation. Following a preoperative MRI, a Leksell stereotactic frame (Elekta AB, Sweden) was applied to the patient’s’ head and a stereotactic computed tomography was obtained. Imaging data was transferred to the iPlan® Stereotaxy software (Brainlab - Feldkirchen, Germany) for calculation of optimal tridimensional coordinates for the inferior and superior aspects of the proposed lesion, as well as for the entry point and trajectory of the radiofrequency probe. A two millimeter straight probe was introduced though a twist drill burr hole in the coordinates of the inferior aspect of the lesion, medial part. A radio frequency lesion was then applied at 80°C for 90 seconds. The probe was adjusted three millimeters laterally, and a second lesion was placed, followed by a third and fourth lesion at the superior coordinates. This procedure was repeated for the contralateral amygdala. Figure 5 illustrates probe placement.

### MRI AQUISITION

Scans were obtained on a 1.5 Tesla MRI system (Magnetom Espree, Siemens, Germany). T1-weighted structural images (slice thickness 1.0 mm, no gap, TE/TR 5/300 ms, flip angle 45°, FOV 240 mm, 1 x 0.9 x 0.9 mm or 1 x 0.5 x 0.5 mm voxel) and diffusion-weighted images (DWI) (4.0 mm slice thickness, gap 0.5 mm, TE/TR 5000/100 ms, flip angle 45°, FOV 240 mm, 4 x 2 x 2 mm voxel, b=1000, 36 directions, 3 b0 images) were acquired. Images were resampled to 0.5mm isotropic voxels for Deformation Based Morphometry (DBM) analysis. The Fractional Anisotropy (FA) and Mean Diffusivity (MD) maps were computed at native resolution and then coregistered with the resampled T1-weighted images resulting from the DBM analysis.

### IMAGE PROCESSING

Presurgical amygdala volume and post-surgical lesion size were estimated using Amira® 3D (FEI Houston Company, USA). For the remaining analysis, amygdala lesion and area of the probe entry were excluded. T1-weighted images were non-uniformity corrected and skull-stripped^33^. To capture changes in local brain morphology, we used DBM, a technique based on image registration. Image registration produces transformations that map every point in one image to its corresponding point in another image. Thus, it allows for detection and quantification of anatomic differences. In this sense, longitudinal changes were evaluated by DBM using the ANTs toolkit (http://stnava.github.io/ANTs/; ‘antsMultivariateTemplateConstruction2.sh’)^33^ for linear and non-linear registration in a 2 level registration procedure where we first registered all images of the same subject to one another to detect within subject changes occurring over time – the critical variable of interest for the analysis of the lesion effect.

In a second registration, we registered the subject averages to one another to achieve voxel correspondence to compare the local individual changes across the group of subjects. Local volumetric changes of each voxel within each image are captured in the Jacobian of the deformation from the image registration, and have been shown to be sensitive to anatomical changes within the brain without requiring a priori segmentation^25^. Especially within-subject longitudinal DBM has been shown to be capable of detecting small local brain changes^34^. The Jacobians were calculated within each subject, transformed to the group average and smoothed using a 2 mm gaussian 3D kernel.

DWI images were analyzed using the MRtrix package (J-D Tournier, Brain Research Institute, Australia, https://github.com/MRtrix3/mrtrix3)^35^. Raw images were first corrected for distortions and motion^36^ within each acquisition, and then denoised^37^ before calculating the diffusion tensor. FA and MD maps were then computed from the diffusion tensor. To achieve local correspondence between the DWI and the DBM measures, the individual DWI images were coregistered with the corresponding individual T1-weighted images and mapped onto the group average of the DBM analysis. FA and MD maps were subsequently blurred using a 2 mm gaussian 3D kernel. Local MD and FA values were extracted in regions that showed significant local volume changes to investigate the possible microstructural changes.

### STATISTICS

The RMINC (https://github.com/Mouse-Imaging-Centre/RMINC) package in RStudio (version 3.3.3, https://www.r-project.org/) was used for statistical analysis. A linear mixed-effect model was performed. To test changes over time and the relationship between behavior and hormone evaluation we used a linear mixed effect model with subject as random effect.

For imaging analysis, the model tested local change at every voxel over time with the three timepoints as ordered factor as fixed and patient and random effect. T-statistics for linear and quadratic trajectories of change across time were calculated at each voxel and corrected for multiple comparisons using false discovery rate (FDR). The level of significance was set at q<0.05 (whole brain FDR-correction) for all DBM analyses. To investigate possible patterns in the DBM results, a hierarchical clustering analysis was performed. The input was a cross correlation matrix, where element (x,y) describes the correlation of computed Jacobians between region x and region y across all 4 subjects. The agglomerative clustering algorithm from sci-kit learn (http://scikit-learn.org/) was used with ward linkage criterion. In this method, each observation (i.e., vector of correlations) starts as its own cluster before clusters are successively merged together based on their similarity.

Maps of volumetric change were used to identify regions of interest (ROI) for inside/outside genetic enrichment analysis of areas that grew or shrunk significantly over time compared to the rest of the brain using the Allen Institute human gene expression data^38^ (© 2010 Allen Institute for Brain Science. Allen Human Brain Atlas. human.brain-map.org).T-stat maps were thresholded at |t| >2.66 (q<0.1 whole brain FDR corrected; this threshold is necessary to get a more expansive spatial map of volumetric changes to colocalize with brain areas included in the Allen brain atlas). The well locations used for the Allen gene atlas were registered to common space and each well location was given a spatial extent of a sphere with radius 2mm. T-stat maps were registered to common space, binarized, and a mixed effects linear model with Age and Sex as fixed effect covariates as well as Allen subject ID as a random effect covariate were used to compare the genetic expression inside versus outside the binarized map. Inside versus outside gene significance was then adjusted for multiple comparisons using bonferroni correction (p_cor_<0.05) sorted and split for positive and negative t-statistics (genes enriched versus underexpressed within versus the rest of the brain). Finally the ranked gene orders were used as inputs for gene enrichment analysis using the Gorilla^39^ and GEAS^40^ tools.

## ACKNOWLEDGEMENTS

The authors thanks all the research assistants and staff of Hospital Sirio-Libanes and HCFMUSP. RCRM and FVG are the recipients of grants FAPESP #11/08575-7, #13/20602-5 and #17/10466-8 from Brazil’s government. HB is recipient of grants FAPESP and CNPQ.

## DISCLOSURES

The authors report no conflicts of interest.ch7ng 2001

---

**Supplementary Figure 1.**
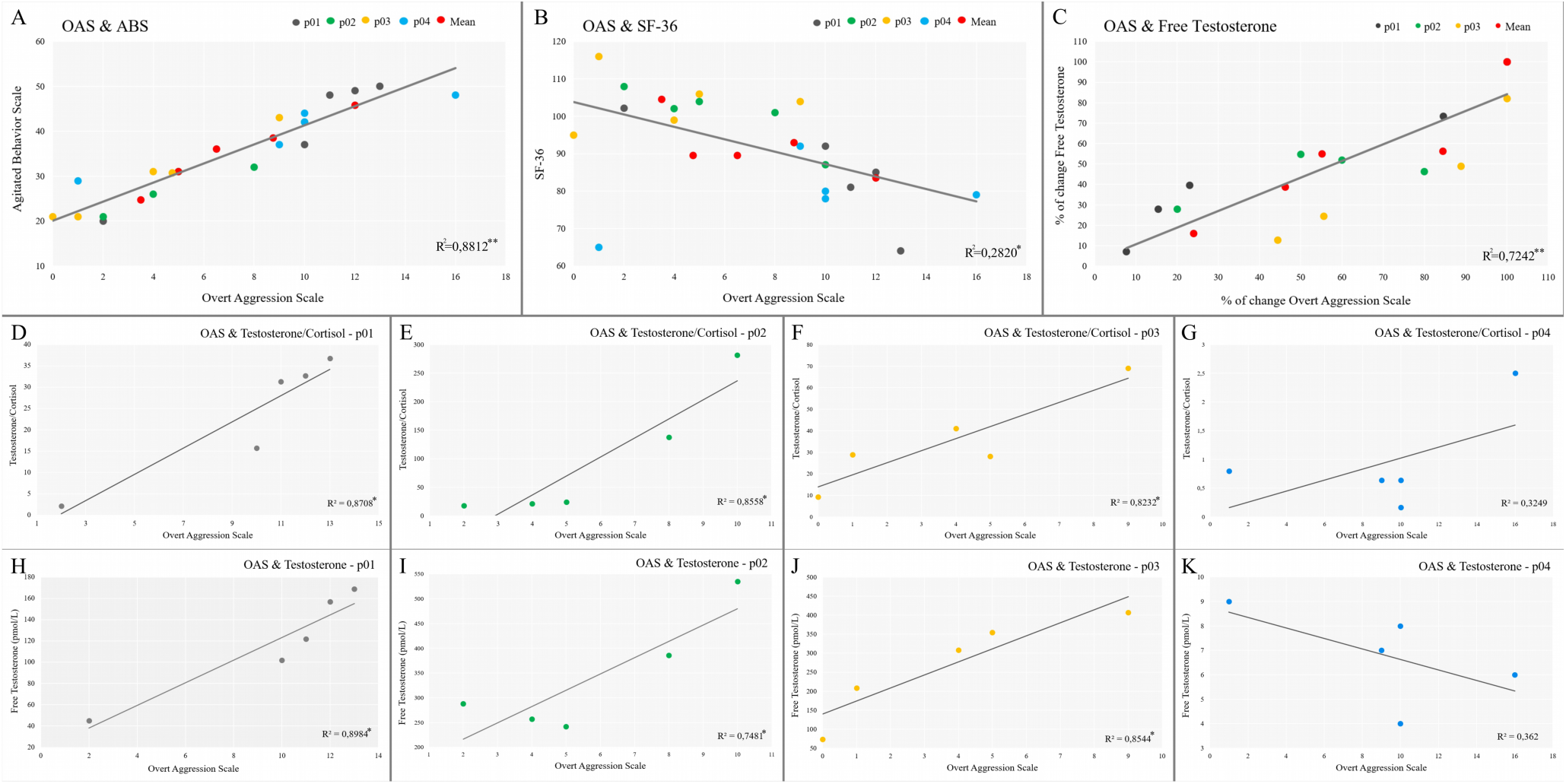
Hormonal and behavior correlations. A. Aggressive Behavior (Overt Aggression Scale - OAS) and Agitation (Agitated Behavior Scale - ABS). B. Aggressive Behavior (OAS) and Quality of life (SF-36). C. % of change in aggressive behavior (OAS) and % of change in Free Testosterone. D-G. Individual analysis of aggressive behavior and free testosterone (p01-p04). H-K. Individual analysis of aggressive behavior and testosterone/cortisol ratio (p01-p04).*p<0.05; **p<0.01

**Supplementary Figure 2.**
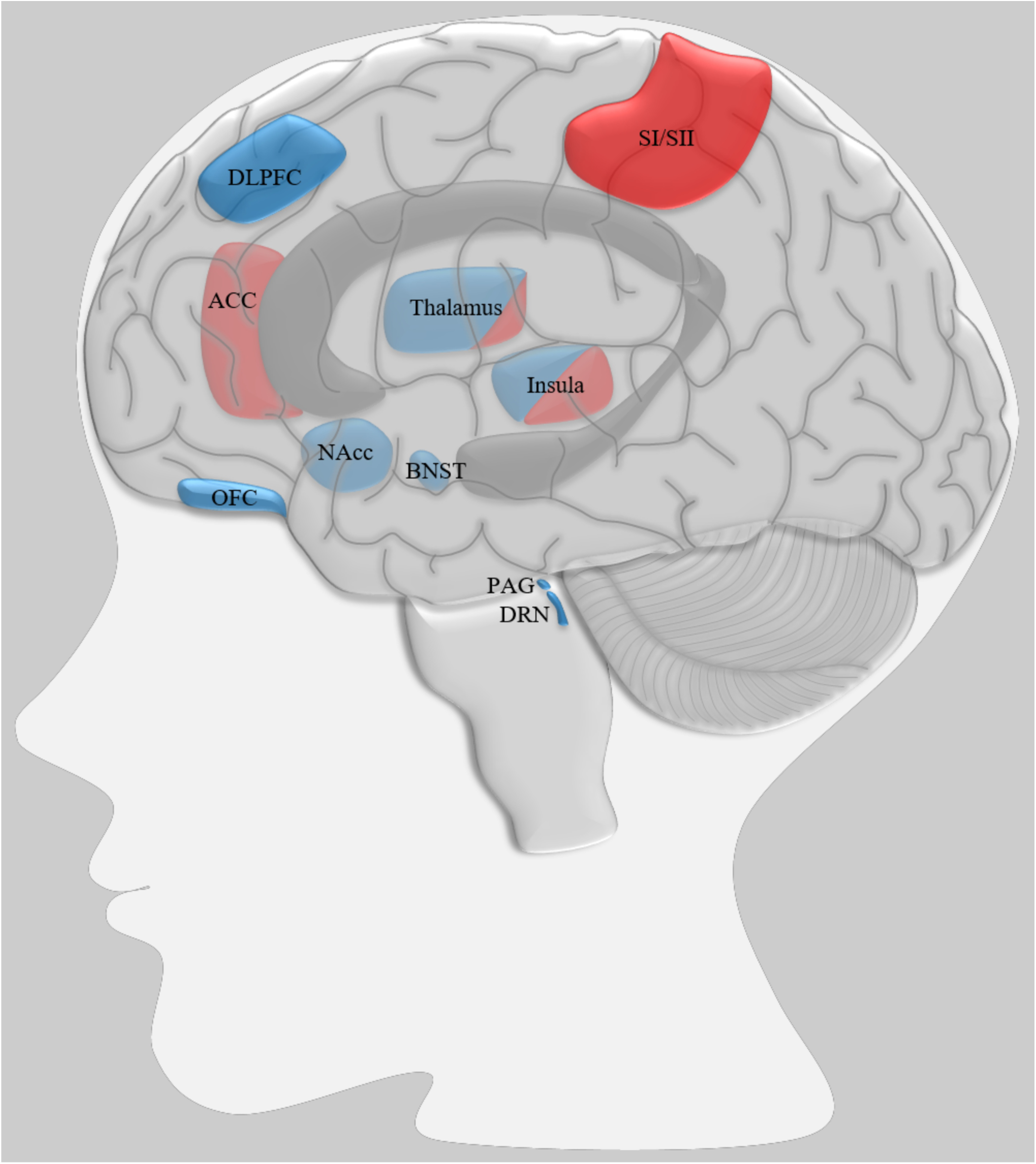
Illustration representing the main changes in areas related to cluster 1 aggressive behavior (blue, decreases in volume) and cluster 2 somatosensation (red, increases in volume). Abbreviations: ACC: anterior cingulate cortex; BNST: bed nucleus of stria terminalis; DLPFC: dorsolateral prefrontal cortex; DRN: dorsal raphe nucleus; NAcc: nucleus accumbens; OFC: orbitofrontal cortex; PAG: periaqueductal gray; SI/SII: somatosensory cortex.

**Supplementary Figure 3:**
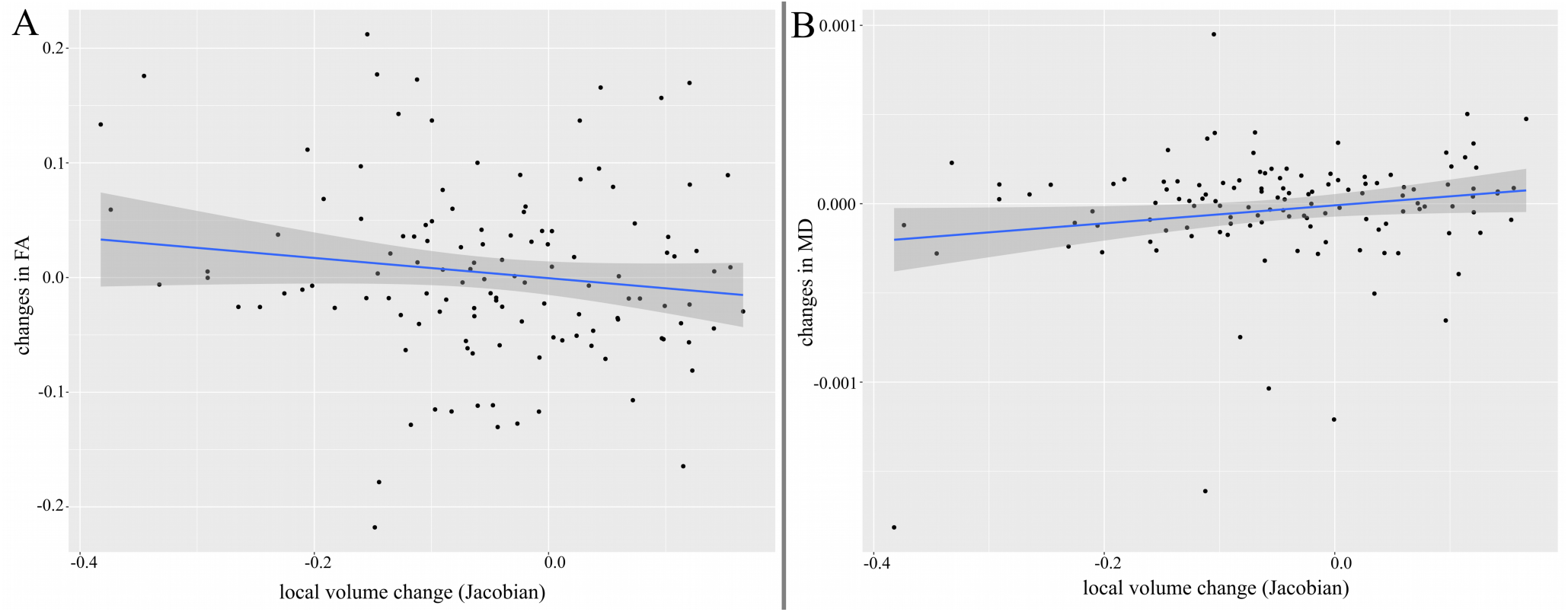
Correlation between DBM Jacobians, FA and MD maps for all peaks. A. Correlations between local volume change in Jacobian and FA map. B. Correlations between local volume change in Jacobian and MD map.

**Supplementary Figure 4.**
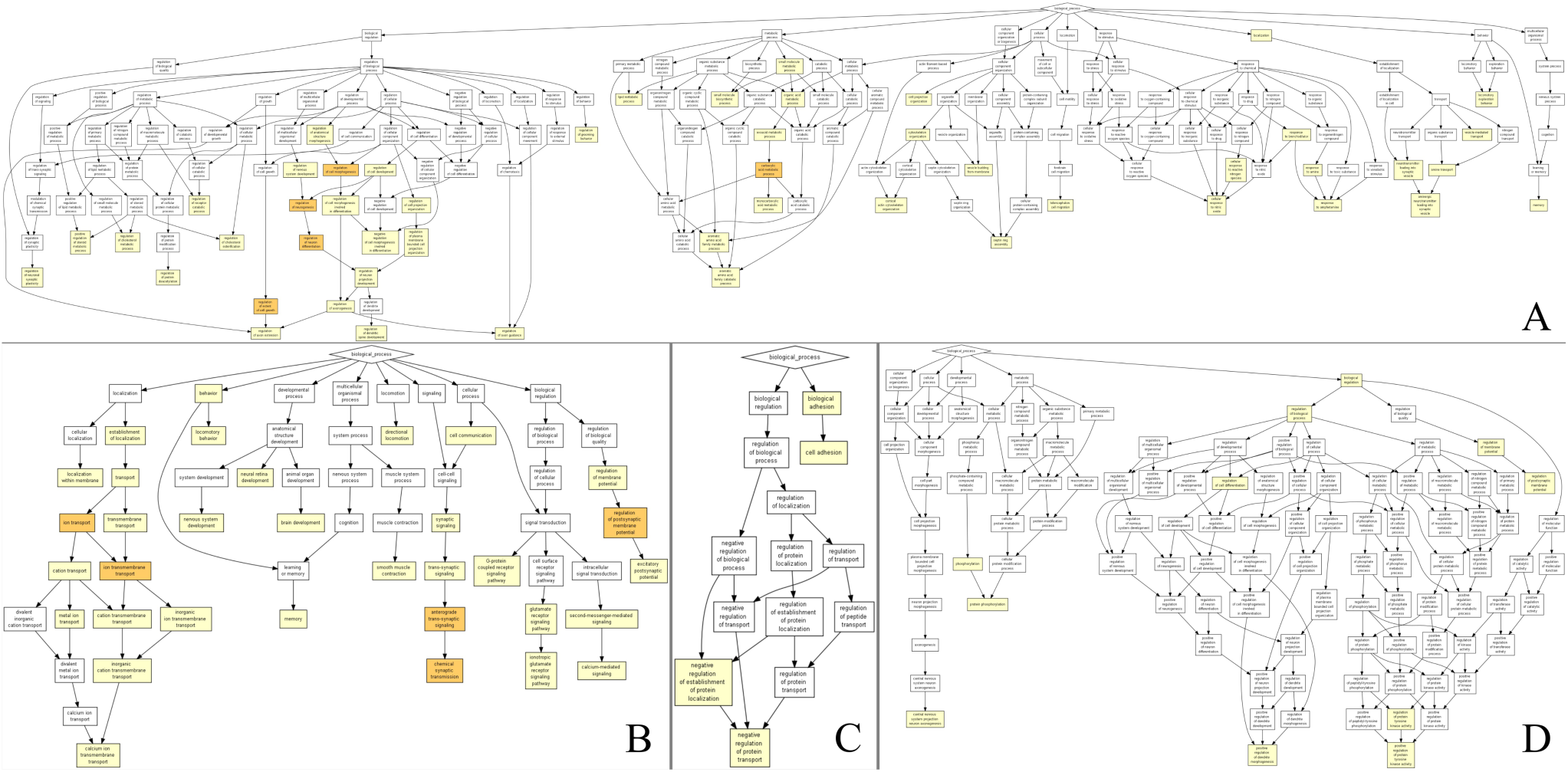
Results of gene enrichment analysis using the GOrilla gene ontology tool. A. Highly expressed genes in areas of volumes decrease (DBM). B. Low expressed genes in areas of volumes decrease (DBM). C. Low expressed genes in areas of volumes increase (DBM). D. Highly expressed genes in areas of volumes increase (DBM).

**Supplementary Figure 5.**
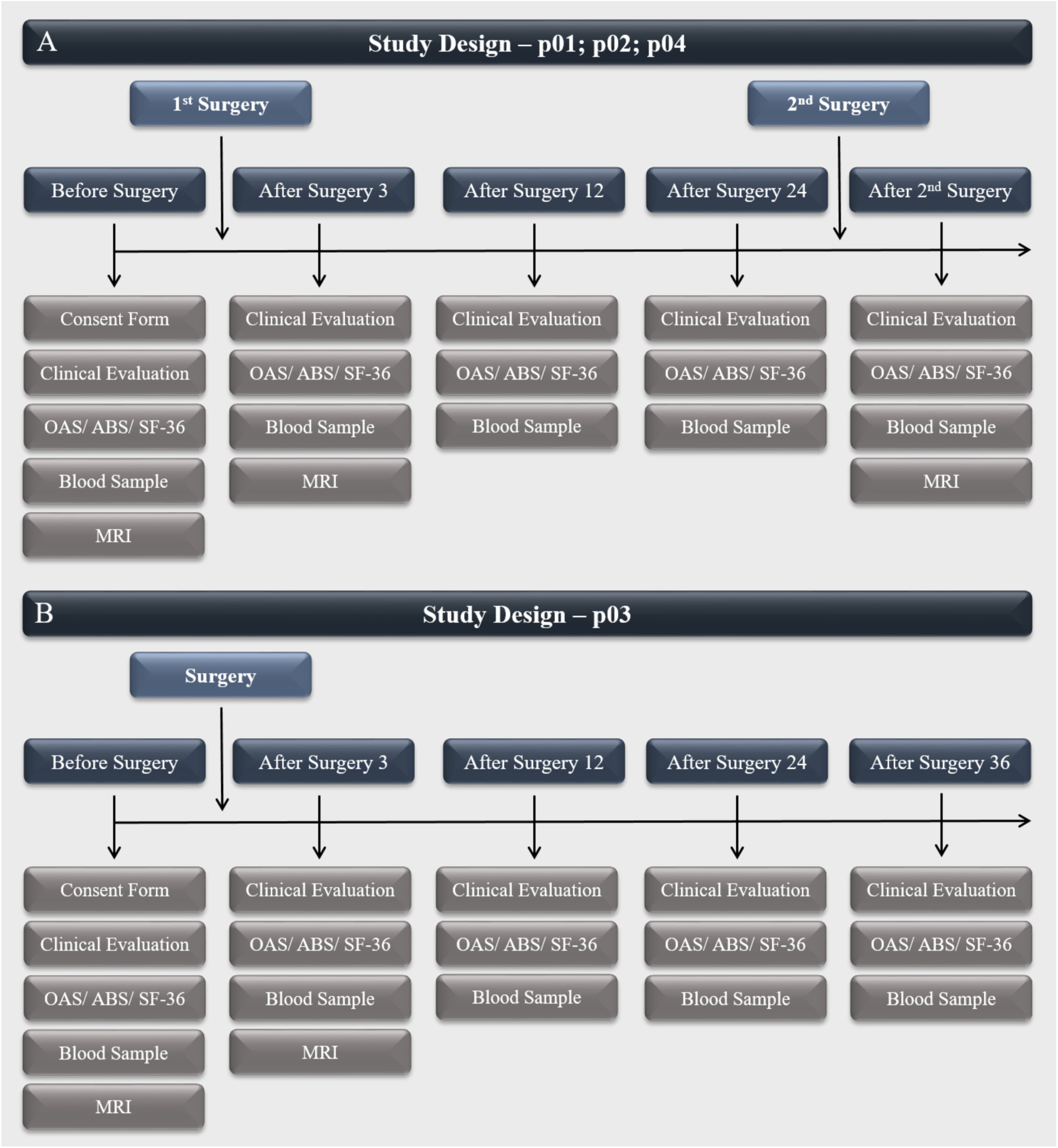
Study design. A. Study design for patients p01, p02 and p04. B. Study design for patient p03. Abbreviations: After surgery 3, 12, 24 and 36: 3, 12, 24 and 36 months after surgery, respectively; OAS: Overt Aggression Scale; ABS: Agitated Behavior Scale; MRI: Magnetic Resonance Imaging.

**Supplementary Table 1.** Data from laboratorial blood test.

|  | Patient p01 |  |  |  |  | Patient p02 |  |  |  |  | Patient p03 |  |  |  |  | Patient p04 |  |  |  |  |
| --- | --- | --- | --- | --- | --- | --- | --- | --- | --- | --- | --- | --- | --- | --- | --- | --- | --- | --- | --- | --- |
|  | 1<sup>st</sup> | 2<sup>nd</sup> | 3<sup>rd</sup> | 4<sup>th</sup> | 5<sup>th</sup> | 1<sup>st</sup> | 2<sup>nd</sup> | 3<sup>rd</sup> | 4<sup>th</sup> | 5<sup>th</sup> | 1<sup>st</sup> | 2<sup>nd</sup> | 3<sup>rd</sup> | 4<sup>th</sup> | 5<sup>th</sup> | 1<sup>st</sup> | 2<sup>nd</sup> | 3<sup>rd</sup> | 4<sup>th</sup> | 5<sup>th</sup> |
| TSH (mIU/L) | 1.83 | 1.58 | 2.24 | 2.16 | 2.76 | 0.89 | 0.91 | 1.11 | 0.94 | 1.74 | 1.75 | 0.78 | 1.39 | 1.49 | 2.32 | 2.56 | 1.56 | 1.99 | 3.66 | 1.13 |
| Free T4 (ng/dL) | 1.58 | 1.18 | 1.01 | 1.25 | 01.09 | 1.44 | 1.48 | 1.49 | 1.48 | 1.72 | 1.16 | 0.94 | 0.75 | 0.91 | 0.84 | 1.31 | 01.04 | 1.33 | 0.99 | 1.16 |
| T3 (mg/dL) | 0.13 | 0.10 | 0.10 | 0.10 | 0.12 | 110 | 110 | 120 | 120 | 0.15 | 0.81 | 0.81 | 0.72 | 0.69 | 0.81 | 0.13 | 0.07 | 0.09 | 0.10 | 0.09 |
| Cortisol (mg/dL) | 4.60 | 3.90 | 6.50 | 4.80 | 21.50 | 1.90 | 10.10 | 12.20 | 2.80 | 16.5 | 5.90 | 7.90 | 7.50 | 12.60 | 7.20 | 2.40 | 12.50 | 11.30 | 24.16 | 11.00 |
| LH (IU/L) | 1.70 | 6.00 | 1.40 | 1.30 | 2.30 | 3.40 | 0.80 | 9.50 | 6.30 | 8.4 | 1.80 | 2.00 | 7.00 | 10.40 | 5.60 | 6.30 | 0.30 | 18.80 | 9.98 | 14.50 |
| FSH (IU/L) | 1.30 | 2.40 | 0.60 | 0.90 | 0.60 | 5.90 | 3.60 | 7.40 | 6.80 | 3.8 | 2.00 | 1.70 | 2.90 | 3.00 | 3.10 | 5.10 | 2.10 | 6.20 | 6.61 | 5.80 |
| Estradiol (pg/mL) | 17.00 | 19.00 | 15.00 | 21.10 | 24.80 | 55.00 | 23.00 | 24.10 | 44.30 | 30.2 | 30.00 | 18.00 | 15.00 | 15.00 | 29.50 | 27.00 | 16.00 | 18.50 | 26.79 | 15.00 |
| Prolactin (mg/dL) | 0.57 | 0.69 | 0.69 | 0.85 | 0.74 | 3.20 | 4.30 | 5.30 | 5.80 | 4.2 | 60.10 | 19.20 | 7.50 | 8.40 | 11.60 | 95.40 | 45.30 | 107.9 | 100.2 | 64.10 |
| Progesterone (ng/mL) | 0.30 | 0.30 | 0.50 | 0.40 | 0.50 | 0.30 | 0.30 | 0.40 | 0.40 | 0.6 | 0.30 | 0.40 | 0.30 | 0.30 | 0.30 | 0.30 | 0.90 | 0.30 | 0.10 | 0.30 |
| Total Testosterone (ng/mL) | 336 | 198 | 115 | 350 | 115 | 412 | 310 | 281 | 428 | 279 | 409 | 97.00 | 528 | 634 | 359 | 18.00 | 22.00 | 29.00 | 53.73 | 22.00 |
| Free Testosterone (pmol/L) | 169 | 122 | 102 | 157 | 45.00 | 535 | 242 | 257 | 386 | 288 | 407 | 73.00 | 308 | 355 | 208 | 6.00 | 8.00 | 9.00 | 0.40 | 7.00 |
| SHBG (nmol/L) | 57.00 | 45.00 | 50.20 | 62.60 | 76.80 | 17.00 | 20.00 | 26.10 | 21.00 | 13.6 | 26.00 | 26.00 | 44.70 | 52.30 | 48.30 | 81.00 | 78.00 | 93.80 | 113 | 91.90 |
| Urea (mg/dL) | 31.00 | 27.00 | 39.00 | 33.00 | 31.00 | 35.00 | 19.00 | 33.00 | 25.00 | 28.00 | 22.00 | 20.00 | 20.00 | 19.00 | 13.00 | 28.00 | 28.00 | 18.00 | 31.00 | 20.00 |
| Creatinine (mg/dL) | 0.76 | 0.75 | 0.97 | 0.86 | 0.87 | 0.86 | 0.80 | 0.90 | 0.78 | 0.84 | 0.86 | 0.81 | 0.92 | 0.83 | 0.70 | 0.84 | 0.72 | 0.56 | 0.65 | 0.61 |
| Sodium (mEq/L) | 140 | 144 | 141 | 138 | 141 | 140 | 145 | 138 | 145 | 144 | 137 | 130 | 131 | 139 | 129 | 142 | 144 | 142 | 142 | 140 |
| Potassium (mEq/L) | 4.40 | 4.10 | 4.70 | 3.70 | 4.20 | 3.90 | 4.70 | 4.40 | 4.30 | 4.20 | 4.00 | 4.10 | 4.50 | 4.50 | 4.70 | 4.00 | 4.10 | 4.10 | 3.80 | 4.00 |
| AST (U/L) | 19.00 | 21.00 | 23.00 | 11.00 | 16.00 | 16.00 | 16.00 | 12.00 | 16.00 | 14.00 | 25.00 | 21.00 | 17.00 | 19.00 | 21.00 | 25.00 | 27.00 | 29.00 | 32.00 | 29.00 |
| ALT (U/L) | 23.00 | 23.00 | 23.00 | 17.00 | 15.00 | 20.00 | 15.00 | 11.00 | 14.00 | 17.00 | 26.00 | 23.00 | 17.00 | 17.00 | 21.00 | 40.00 | 37.00 | 41.00 | 40.00 | 39.00 |
| Total Bilirubin (mg/dL) | 0.16 | 0.38 | 0.54 | 0.16 | 0.29 | 0.28 | 0.56 | 0.32 | 0.42 | 0.36 | 0.25 | 0.32 | 0.28 | 0.42 | 0.36 | 0.33 | 0.45 | 0.47 | 0.37 | 0.39 |
| Total Cholesterol (mg/dL) | 221 | 221 | 181 | 180 | 227 | 159 | 157 | 142 | 153 | 149 | 256 | 245 | 238 | 261 | 212 | 149 | 137 | 152 | 148 | 163 |
| Triglycerides (mg/dL) | 218 | 103 | 116 | 137 | 208 | 164 | 131 | 138 | 146 | 195 | 183 | 245 | 280 | 373 | 176 | 131 | 131 | 132 | 131 | 130 |
| HDL (mg/dL) | 39.00 | 52.00 | 51.00 | 60.00 | 46.00 | 35.00 | 28.00 | 32.00 | 40.00 | 42.00 | 67.00 | 48.00 | 56.00 | 46.00 | 65.00 | 26.00 | 32.00 | 24.00 | 26.00 | 36.00 |
| LDL (mg/dL) | 138 | 115 | 107 | 93.00 | 139 | 96.00 | 75.00 | 85.00 | 76.00 | 70.00 | 110 | 136 | 123 | 151 | 112 | 73.00 | 75.00 | 82.00 | 77.00 | 79.00 |
| C-reactive protein (mg/L) | 2.60 | 3.20 | 12.60 | 0.90 | 1.30 | 11.20 | 7.60 | 3.10 | 9.60 | 6.30 | 1.00 | 2.30 | 1.50 | 1.70 | 2.00 | 1.60 | 2.30 | 1.80 | 2.20 | 3.00 |
| Platelets (th/mm3) | 301 | 292 | 327 | 212 | 267 | 165 | 135 | 234 | 184 | 196 | 214 | 173 | 183 | 202 | 166 | 96.00 | 89.00 | 92.00 | 102 | 83.00 |
| MPV (fL) | 9.70 | 9.70 | 9.90 | 9.50 | 9.00 | 11.80 | 11.30 | 12.30 | 11.60 | 11.80 | 9.70 | 10.00 | 9.40 | 9.00 | 9.10 | 11.90 | 11.80 | 11.90 | 12.30 | 11.90 |
| Erythrocytes (mil/mm3) | 5.49 | 4.94 | 5.36 | 4.92 | 4.93 | 5.49 | 5.16 | 5.75 | 5.58 | 5.48 | 4.66 | 4.46 | 4.47 | 4.42 | 4.37 | 4.38 | 3.93 | 4.38 | 4.28 | 4.65 |
| Hemoglobin (g/dL) | 16.20 | 14.70 | 16.20 | 15.50 | 15.70 | 13.60 | 13.00 | 12.70 | 12.10 | 14.90 | 14.60 | 14.10 | 14.00 | 14.10 | 13.70 | 14.40 | 12.60 | 14.00 | 13.80 | 14.90 |
| Hematocrit (%) | 46.20 | 42.00 | 46.30 | 44.90 | 44.30 | 41.40 | 39.90 | 40.70 | 39.00 | 44.70 | 41.6 | 39.90 | 40.30 | 38.20 | 38.70 | 42.50 | 36.80 | 40.20 | 40.00 | 42.40 |
| MCV (fL) | 84.20 | 85.00 | 86.40 | 91.30 | 89.90 | 75.40 | 77.30 | 70.80 | 69.90 | 81.60 | 89.30 | 89.50 | 90.20 | 86.40 | 88.60 | 97.00 | 93.60 | 91.80 | 93.50 | 91.20 |
| MCH (pg) | 29.50 | 29.80 | 30.20 | 31.50 | 31.80 | 24.80 | 25.20 | 22.10 | 21.70 | 27.20 | 31.30 | 31.60 | 31.30 | 31.90 | 31.40 | 32.90 | 32.10 | 32.00 | 32.20 | 32.00 |
| MCHC (g/dL) | 35.10 | 35.00 | 35.00 | 34.50 | 35.40 | 32.90 | 32.60 | 31.20 | 31.00 | 33.30 | 35.10 | 35.30 | 34.70 | 36.90 | 35.40 | 33.90 | 34.20 | 34.80 | 34.50 | 35.10 |
| RDW-CV (%) | 12.80 | 12.80 | 12.30 | 12.90 | 12.60 | 19.80 | 19.30 | 20.00 | 19.10 | 16.50 | 12.30 | 12.20 | 12.10 | 12.00 | 12.30 | 12.50 | 12.20 | 12.40 | 12.10 | 12.10 |
| RDW-SD (fL) | 38.80 | 39.00 | 38.60 | 42.20 | 41.30 | 51.70 | 53.10 | 50.00 | 48.70 | 48.70 | 39.50 | 39.00 | 39.30 | 37.00 | 39.80 | 43.60 | 40.90 | 42.00 | 40.90 | 39.50 |
| Leukocytes (th/mm3) | 4.77 | 6.19 | 4.99 | 9.48 | 5.95 | 09.01 | 5.46 | 7.47 | 6.86 | 6.58 | 4.62 | 6.18 | 4.49 | 4.20 | 4.30 | 3.76 | 3.81 | 2.67 | 4.60 | 6.61 |
| Neutrophils (%) | 42.10 | 85.40 | 37.10 | 39.70 | 30.20 | 75.30 | 63.60 | 5.47 | 71.90 | 65.80 | 42.10 | 37.00 | 38.30 | 34.20 | 30.20 | 49.00 | 50.70 | 48.00 | 45.00 | 84.40 |
| Eosinophils (%) | 4.60 | 0.20 | 5.20 | 0.30 | 4.50 | 1.90 | 2.90 | 0.15 | 2.90 | 1.70 | 0.40 | 0.50 | 0.20 | 0.70 | 0.50 | 1.60 | 2.60 | 0.70 | 1.30 | 0.30 |
| Basophils (%) | 0.20 | 0.00 | 0.20 | 0.10 | 0.20 | 0.60 | 0.50 | 0.05 | 0.60 | 0.60 | 0.40 | 0.20 | 0.40 | 0.50 | 0.00 | 0.00 | 0.30 | 0.40 | 0.20 | 0.00 |
| Lymphocyte (%) | 43.00 | 11.80 | 42.70 | 47.50 | 50.80 | 12.80 | 22.70 | 1.19 | 16.30 | 22.30 | 43.50 | 51.90 | 44.80 | 49.80 | 56.70 | 42.80 | 35.40 | 43.80 | 42.00 | 12.90 |
| Monocytes (%) | 10.10 | 2.60 | 14.80 | 12.40 | 14.30 | 9.40 | 10.30 | 0.61 | 8.30 | 9.60 | 13.60 | 10.40 | 16.30 | 14.80 | 12.60 | 6.60 | 11.00 | 7.10 | 11.50 | 2.40 |

Abbreviations: 1<sup>st</sup>: Before surgery; 2<sup>nd</sup>: 3 months after surgery; 3<sup>rd</sup> 12 months after surgery; 4<sup>th</sup> 24 months after surgery; 5<sup>th</sup> 36 months after surgery for p03 and after second surgery for p01, p02, p04; TSH: Thyroid-Stimulating Hormone; Free T4: Free Thyroxine; T3: Triiodothyronine; LH: Luteinizing Hormone; FSH: Follicle-Stimulating Hormone; SHBG: Sex Hormone Binding Globulin; AST: Aspartate Aminotransferase; ALT: Alanine Aminotransferase; HDL: High-Density Lipoprotein; LDL: Low-Density Lipoprotein; MPV: Mean Platelet Volume; MCV: Mean Corpuscular Volume; MCH: Mean Cell Hemoglobin; MCHC: Mean Cell Hemoglobin Concentration; RDW-CV: Red Cell Distribution Width – Cell Volume; RDW-SD: Red Cell Distribution Width – Standard Deviation.

